# Impaired miRNA degradation by post-transcriptional addition of 3’ cytosine and adenine in T cell activation

**DOI:** 10.1101/2020.08.19.257816

**Authors:** Ana Rodríguez-Galán, Sara G Dosil, Manuel José Gómez, Irene Fernández-Delgado, Fátima Sánchez-Cabo, Francisco Sánchez-Madrid

**Affiliations:** Servicio de Inmunología. Hospital Universitario La Princesa, Instituto Investigación Sanitaria Princesa (IIS-IP), Universidad Autónoma de Madrid (UAM), Madrid, Spain; Vascular Pathophysiology Area. Centro Nacional de Investigaciones Cardiovasculares (CNIC), Madrid, Spain; CIBER de Enfermedades Cardiovasculares, Instituto de Salud Carlos III, Madrid, Spain

**Keywords:** miRNA, post-transcriptional modifications, cytosylation, adenylation, T cell activation

## Abstract

MiRNA repertoire of T cells undergoes extensive changes in response to activation. Whereas global miRNA downregulation occurs few hours after activation, some individual miRNAs are specifically up- or down-regulated. In this study, we have assessed miRNA expression and post-transcriptional modification kinetics in human primary CD4+ T cells upon short-term stimulation with αCD3αCD28 or IFN I using Next Generation Sequencing. Multiple miRNAs not related before with T cell activation profile have been identified as differentially expressed. Downregulated miRNAs presented higher 3’ uridylation. Dis3L2 and Eri1 (3’ to 5’ exoribonucleases that prefer uridylated RNA as substrates) increased their expression upon TCR stimulation, probably generating an adverse environment for miRNAs. Remarkably, non-templated cytosine additions to 3’ end, previously unknown to be a relevant post-transcriptional modification mechanism, were overrepresented in upregulated miRNAs, together with high levels of adenylation. In the midst of an increasing presence of exoribonucleases, miRNAs multiplying their levels may successfully escape degradation due to 3’ cytosine and adenine addition. These protective signals open a new avenue to improve miRNA stability for therapy in T cells.

## INTRODUCTION

MiRNA are key modulators that fine-tune immune responses (Mehta and Baltimore 2016; Gracias and Katsikis 2011; Lindsay 2008; Podshivalova and Salomon 2013). During T cell activation, miRNA profile undergoes extensive changes, with a global downregulation of total miRNA levels occurring as early as 4 hours after activation (Bronevetsky et al. 2013). Beyond the overall picture of general reduction, some individual miRNAs stand out for their specific up- or downregulation, as shown by arrays, RT-qPCR and Northern Blot (Bronevetsky et al. 2013; Gutiérrez-Vázquez et al. 2017; Jindra et al. 2010; Wu et al. 2007; Grigoryev et al. 2011; Sousa et al. 2017; Teteloshvili et al. 2015). These studies, which evaluate mouse samples 18 h to 7 d; and human samples 2 to 7 d post-activation have been gathered together in a recent review (Rodríguez-Galán et al. 2018).

Little is known regarding the mechanisms that underlie these changes in the T cell miRNA landscape. Previous research from our laboratory pointed to 3’ uridine addition as a potential mechanism guiding miRNA turnover during T cell activation (Gutiérrez-Vázquez et al. 2017). Uridylation is a relatively common post-transcriptional miRNA modification. Next Generation Sequencing (NGS) has identified not only nucleotide additions to the expected genomic miRNA sequences, but also trimmings and substitutions (Ebhardt et al. 2009; Lee et al. 2010). Post-transcriptional Modifications (PtMs) generate multiple variants of the same miRNA (isomiRs) that differ in their 5’, 3’ or internal modifications. PtMs modulate biogenesis, stability and function (Neilsen et al. 2012; Gebert and MacRae 2019; de Sousa et al. 2019). Several mechanisms elicit PtMs on the canonical miRNA sequence including: alternative processing by Drosha or Dicer, RNA editing and non-template nucleotide addition.

During miRNA biogenesis, Drosha cleaves the primary-miRNA (pri-miRNA) transcript, generating a hairpin precursor-miRNA (pre-miRNA) which is subsequently processed by Dicer, leading to the generation of a double-stranded miRNA duplex. Drosha and Dicer excisions are slightly flexible, thereby becoming a source of 5’ and 3’ isomiRs (Wu et al. 2009; Starega-Roslan et al. 2011; Zhou et al. 2012; Gu et al. 2012; Kim et al. 2017; Zhu et al. 2018; Kwon et al. 2019).

Other forms of PtMs derive from RNA editing which include conversion of adenosine (A) to inosine (I) by ADARs (adenosine deaminases acting on RNA) (Yang et al. 2006; Bazak et al. 2014; Nishikura 2016; Tan et al. 2017); or deamination of cytidine (C) to uridine (U) by APOBECs (apolipoprotein B mRNA editing enzyme, catalytic polypeptide-like) (Blanc and Davidson 2010; Rosenberg et al. 2011). Since I is a guanosine (G) analog, A-to-I editing is equivalent to an A-to-G mutation. For miRNAs, A-to-I editing is well-characterized (Li et al. 2018a; Wang and Liang 2018), whereas the physiological relevance of C-to-U modification is currently unknown.

Additional enzymes responsible for miRNA PtMs are TErminal Nucleotidyl Transferases (TENTs). TENTs catalyze non-template additions (NTAs) of nucleotides mainly at the 3’ end (‘tailing’) (Warkocki et al. 2018). TENTs are often flexible substrate-wise, but those with a preference towards adding adenosine are known as non-canonical poly(A) polymerases (ncPAPs). Other TENTs prefer to add uridine, namely terminal uridyl transferases (TUTases). Uridylation and adenylation are the most typical 3’ end modifications across animal miRNAs (Landgraf et al. 2007; Chiang et al. 2010; Burroughs et al. 2010; Wyman et al. 2011; Muller et al. 2014). Multiple studies have explored these modifications and their consequences in detail. Conclusions often appear contradictory, which likely result from the specific biological context, including different species, cell type or cellular compartment. For instance, GLD-2 (PAPD4/TENT2) 3’ monoadenylation seems to stabilize specific miRNA populations in human fibroblasts (D’Ambrogio et al. 2012) and miRNA-122 in the liver (Katoh et al. 2009). In mouse early embryos, 3’ mono- and oligoadenylation appears to protect certain miRNAs in a context of large degradation (Yang et al. 2016). However, PAPD5 (TENT4B/GLD4/TUT3) adenylates miR-21-5p on 3’, promoting its degradation by PARN (Poly(A)-specific ribonuclease) (Boele et al. 2014). In human monocytes, knocking down PAPD4 showed no overall effect of 3’ adenylation on miRNA stability, but adenylation instead altered miRNA effectiveness through reduction of incorporation into the RNA-induced silencing complex (RISC, the complex where miRNAs induce mRNA degradation or inhibit their translation) (Burroughs et al. 2010). Uridylation also promotes diverse outcomes on miRNA. Pre-let-7 miRNA can be uridylated on 3’ by TUT4 (ZCCHC11) or TUT7 (ZCCHC6) (Heo et al. 2009; Thornton et al. 2012). Lin28 (a RNA-binding protein) binds to pre-let-7 and favors oligouridylation (10-20 uridines), which inhibits subsequent Dicer processing and serves as a signal for Dis3L2 degradation (Heo et al. 2008, 2009; Thornton et al. 2012; Ustianenko et al. 2013; Chang et al. 2013). In the absence of Lin28, pre-let-7 undergoes monouridylation to pursue its maturation process (Heo et al. 2012). Let-7 promotes cell differentiation, and the regulatory mechanism triggered by Lin28 expression maintains pluripotency in stem cells (Heo et al. 2009; Büssing et al. 2008). Non-templated uridine addition also occurs on mature miRNA, such as miR-26, which has been described to undergo 3’-uridylation by ZCCHC11 (TUT4) (Jones et al. 2009). MiR-26a and miR-26b uridylation has been shown to reduce their ability to repress IL-6 (Jones et al. 2009). In addition to Dis3L2, Eri1 is also a 3’ to 5’ exonuclease that prefers uridylated RNA substrates (Hoefig et al. 2013).

In order to gain a mechanistic insight in the early changes occurring at the level of miRNA post-transcriptional modifications, we have studied the effect of stimulation of human primary CD4 T cells on miRNA through Next Generation Sequencing (NGS). Since large changes in total miRNA levels occur very early (Bronevetsky et al. 2013), 3 and 6 h were chosen as intermediate points for our time course.

## RESULTS

### 1. MiRNA modulation by αCD3αCD28

A total of 120 miRNAs were differentially expressed (adjusted p value <0.1, 62 upregulated and 58 downregulated) upon stimulation of resting human CD4 T cells with αCD3αCD28 for 3, 6 and 24 h [Fig. 1A,1B]. The most upregulated miRNAs (fold change indicated in brackets) at 24 h were miR-4455 (122x), miR-222-5p (24x), miR-7974 (21x), miR-155-3p (18x) and miR-4521 (15x) [Fig. 1B]. Remarkably, miR-1281 showed a 14 fold change upregulation at 3 h which was maintained at 6 h, but vanished at 24 h. The most downregulated miRNAs at 24h were miR-4485-3p (−11x), miR-4695-3p (−5x), miR-570-3p (−4x) and miR-1260a (−4x). With the exception of miR-155-3p, none of these miRNAs had been connected previously to T cell activation, probably due to the lack of studies using an unbiased approach such as NGS. Consistent with previous evidence (Rodríguez-Galán et al. 2018), we also found downregulation of miR-150 and miR-223; while miR-155, miR-17-5p and miR-18a-5p were upregulated. Ingenuity pathway analysis (IPA) indicated that differentially expressed miRNAs were mainly involved in processes related to cell development, growth, proliferation and movement [Supplementary Fig. 1A]. Networks of predicted targets for the miRNAs with the highest up- and down-regulation show large overlapping with 59 genes targeted by at least 2 of the 6 most upregulated miRNAs and 149, targeted by at least 2 of the 6 most downregulated [Supplementary Fig. 2A].

**Figure 1.**
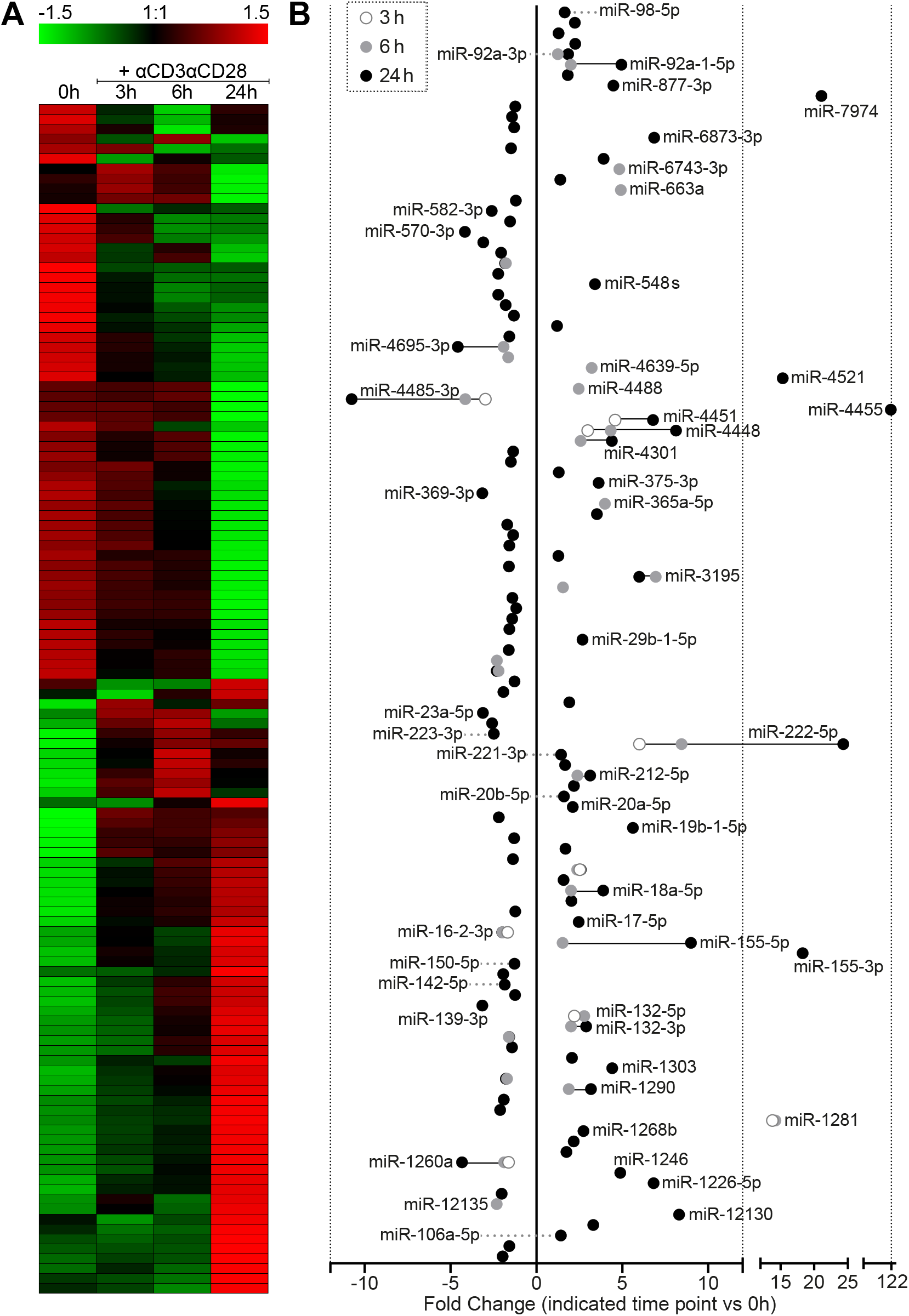
Differential miRNA expression 3h, 6h and 24h after αCD3αCD28 stimulation of human primary CD4+ T cells. A) Heat map for differentially expressed miRNAs. The heatmap represents relative expression values for a non-redundant collection of differentially expressed miRNAs (adjusted p-value <0.1), detected after stimulation with αCD3αCD28 for 3h, 6h, and 24h, relative to 0h. B) Log fold change of differentially expressed miRNAs at 3, 6 and 24h compared to 0h. Representative miRNAs names are included, particularly those with higher fold changes.

### 2. MiRNA modulation by IFN I

In a separate set, resting human CD4+ T cells were stimulated with IFN type I. IFN I significantly altered the expression levels of 57 miRNAs (adjusted p value <0.1): 24 microRNAs were upregulated, and 33 were downregulated [Fig. 2A, 2B]. Compared with the data in Fig. 1, we found that 37 miRNAs were common to the IFN I and αCD3αCD28 subsets (15 upregulated and 21 downregulated in both stimulations, and 1 miRNA regulated in opposite directions) [Fig. 2C]. The most upregulated miRNAs (fold change indicated in brackets) at 24 h were miR-1281 (15x, 3h), miR-3195 (7x, 6h) and miR-3614-5p (8x, 24h). The most downregulated miRNAs at 24h were miR-27a-5p (−14x) and miR-4485 (−15x) [Fig. 2A, 2B]. IPA revealed that most processes controlled by IFN I-regulated miRNAs were very similar to those observed for cells stimulated through αCD3αCD28, mainly: cell development, movement, growth and proliferation [Supplementary Fig. 1B]. Predicted targets show a more intense network overlapping among the 6 most downregulated miRNA with 189 targets common to at least 2 miRNAs; while 52 genes would be targeted by at least 2 of the six most upregulated [Supplementary Fig. 2B].

**Figure 2.**
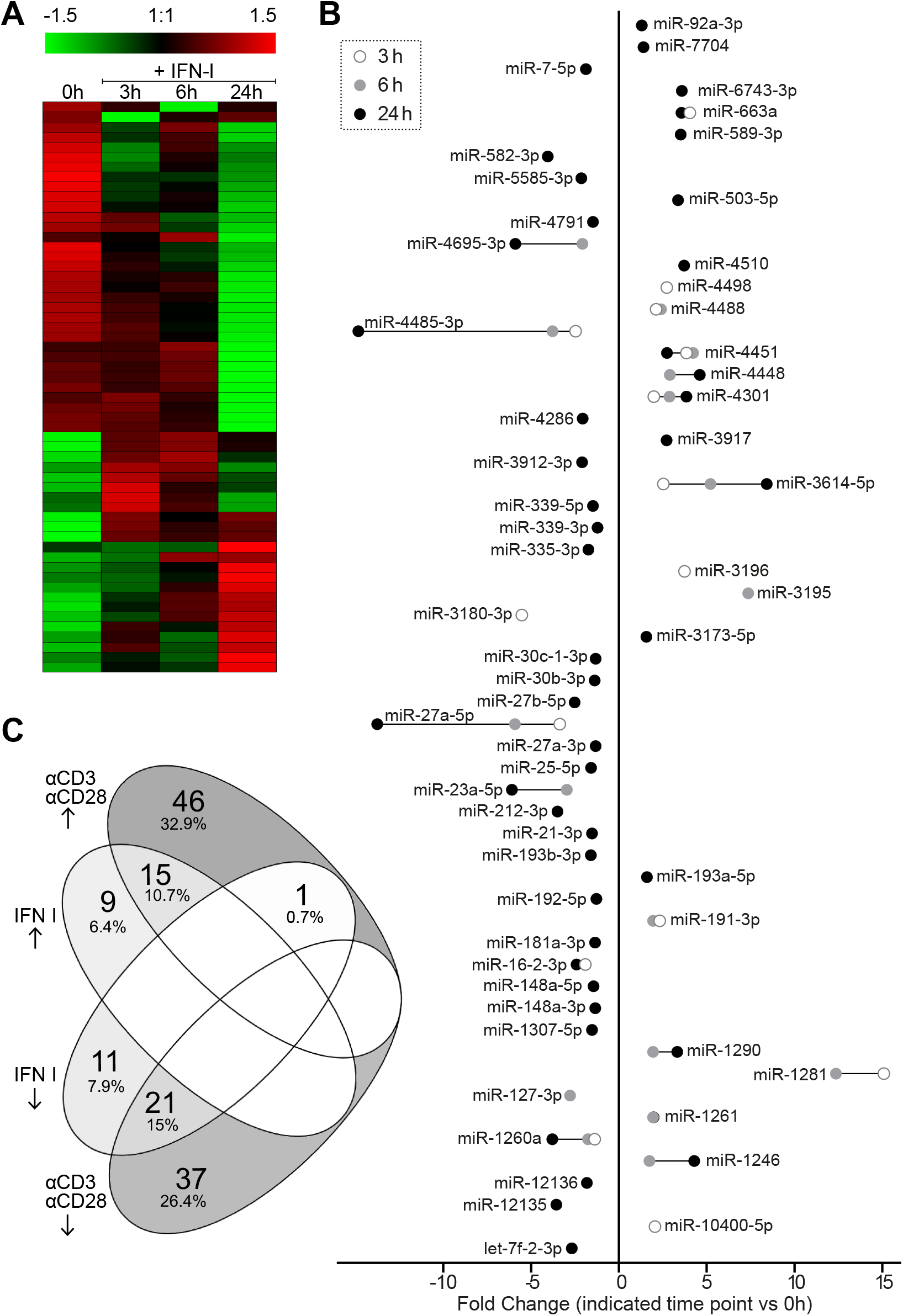
Differential miRNA expression 3h, 6h and 24h after IFN I stimulation of human primary CD4+ T cells. A) Heat map for differentially expressed miRNAs. The heatmap represents relative expression values for a non-redundant collection of differentially expressed miRNAs (adjusted p-value <0.1), detected after stimulation with IFN I for 3h, 6h, and 24h, relative to 0h. B) Log fold change of differentially expressed miRNA at 3, 6 and 24h compared to 0h. C) Venn diagram with miRNAs differentially up and down-regulated by IFN I and αCD3αCD28.

### 3. Post-transcriptional modifications (PtMs)

A global assessment of PtMs indicated that miRNAs in our samples underwent extensive 3’-end modification compared to their canonical sequences [Fig. 3A]. Unexpectedly, C addition was highly represented in our samples at the most modified position: 0 or ‘3p end nucleotide’. A and C modifications were similarly represented in this position, much more frequently than U and G [Fig. 3A]. Modifications at 5’ end and ADAR editing (A to G) were detected on a very limited basis. According to the global profile, positions 0 (3p end) and +1 (3p end +1), were by far the most heavily modified, followed by positions −1 and +2 [Fig. 3A]. For this reason, nucleotide modifications at the 3’ end were analyzed in greater detail in an attempt to discover specific sequences that could guide miRNA dynamics in T cells. PtMs patterns found in the 3’ end (positions −4 to 4) were evaluated (data not shown), indicating that the most common modifications across the different samples were: C, A, U and G mono-additions, and A and U oligo-additions. ‘AU’ was the most frequent multi-nucleotide modification, although sequences combining more than one nucleotide were clearly underrepresented. We also detected UAGU modifications at position −4, as well as AGU and AGUU at −3. We evaluated the presence of U, A, C, G and of UU+, AA+, CC+, GG+ (homopolymers of two or more equal nucleotides) at highly modified 3’-end positions. The results were conjoined for the different stimulation time points [Fig. 3B]. Analyzed PtMs remained stable during activation and were clearly associated to a specific 3’ end position [Fig. 3B].

**Figure 3.**
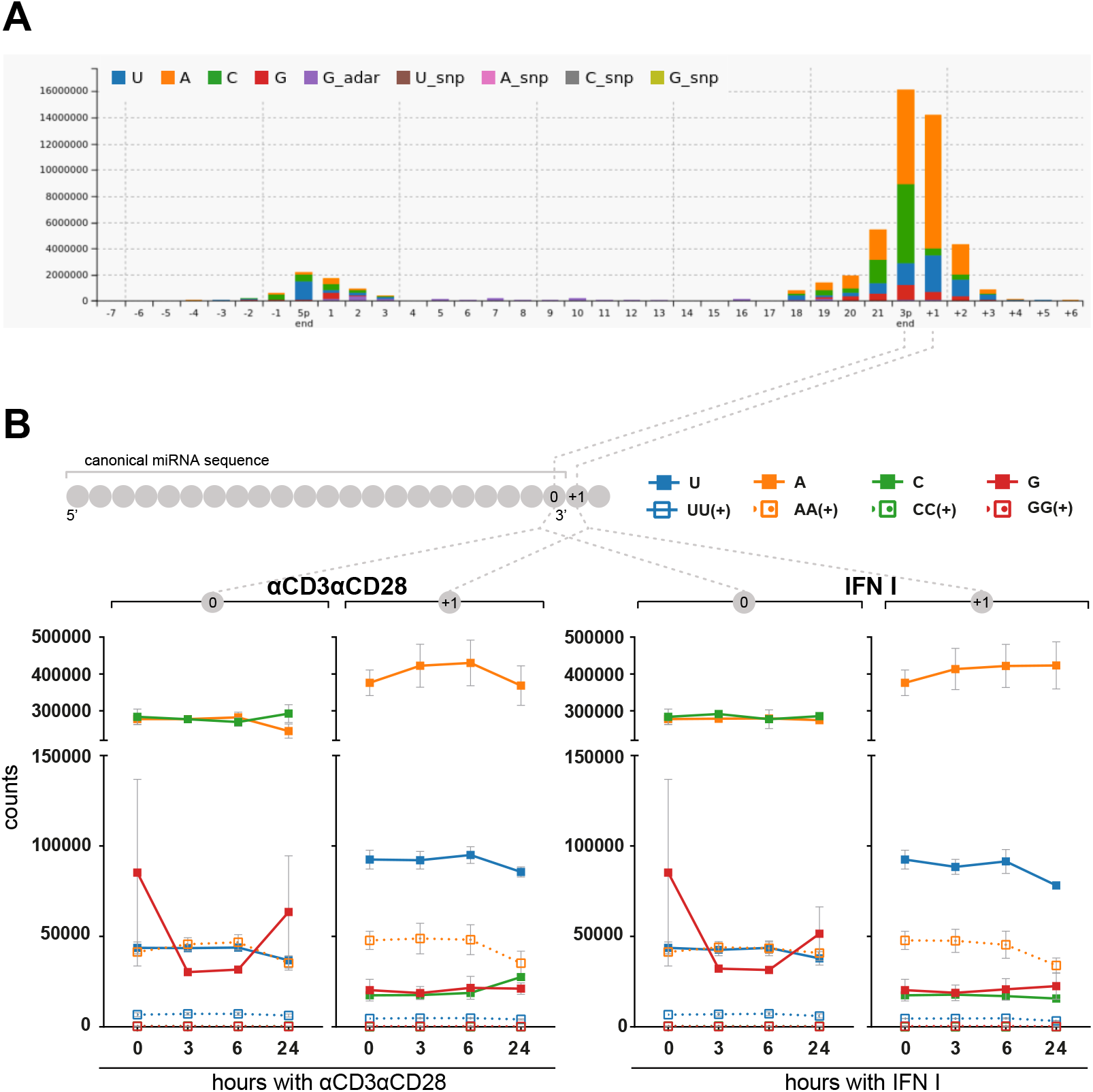
Post-transcriptional miRNA modifications: a global view. A) Global post-transcriptional modifications (PtMs) profile for all 21 sequenced samples generated by Chimira. B) Kinetics of most abundant PtMs (mono-additions: U, A, C, G; and oligo-additions: ≥UU, ≥AA, ≥CC, ≥GG) at positions ‘3p-end’ (0) and ‘3p-end + 1’ (+1), in the population of 626 expressed miRNA species in unstimulated conditions and during activation with αCD3αCD28 (left) or IFN I (right). Mono-additions refer to the specific nucleotide on their own or followed by a different nucleotide, but not followed by the same nucleotide. Oligo-additions include PtMs with two or more equal nucleotides.

To assess whether PtMs could be guiding the differential miRNA expression described in Fig. 1 and Fig. 2, modifications at positions −1, 0, +1 and +2, were represented considering only data from miRNA upregulated or downregulated, either upon αCD3αCD28 stimulation [Fig. 4A, 4B] or IFN I [Fig. 5A, 5B]. Data divided according to differential regulation, revealed the presence of a specific ‘PtMs barcode’ for each population. Upregulated miRNAs were more extensively modified, with the distinct signature of high levels of A addition at +1 and C addition at 0 [Fig. 4A, 4B, Fig. 5A, 5B]. αCD3αCD28 downregulated miRNAs show reduced levels of these specific modifications and a marked presence of U additions, mostly at +1 [Fig. 4A, 4B]. Adenine additions at position +1 were much higher in upregulated miRNAs with counts of A mono-additions around 140000-210000, while downregulated miRNAs counts did not go beyond 30000 in αCD3αCD28 stimulation [Fig. 4A] or 2000 in IFN I [Fig. 5A]. Additions of two or more adenines were also a specific signature of upregulated miRNAs at 0 and +1 [Fig. 4A, Fig. 5A]. Strikingly, miRNAs found to be upregulated showed around 20000 cytosine counts at position 0 before stimulation [Fig. 4A, Fig. 5A], which remain stable upon IFN I stimulation [Fig. 5A] and grew up to around 50000 counts after 24h of αCD3αCD28 stimulation [Fig. 4A], whereas downregulated miRNAs maintained their cytosine counts around 5000 and below across all time points [Fig. 4A, Fig. 5A; position 0].

**Figure 4.**
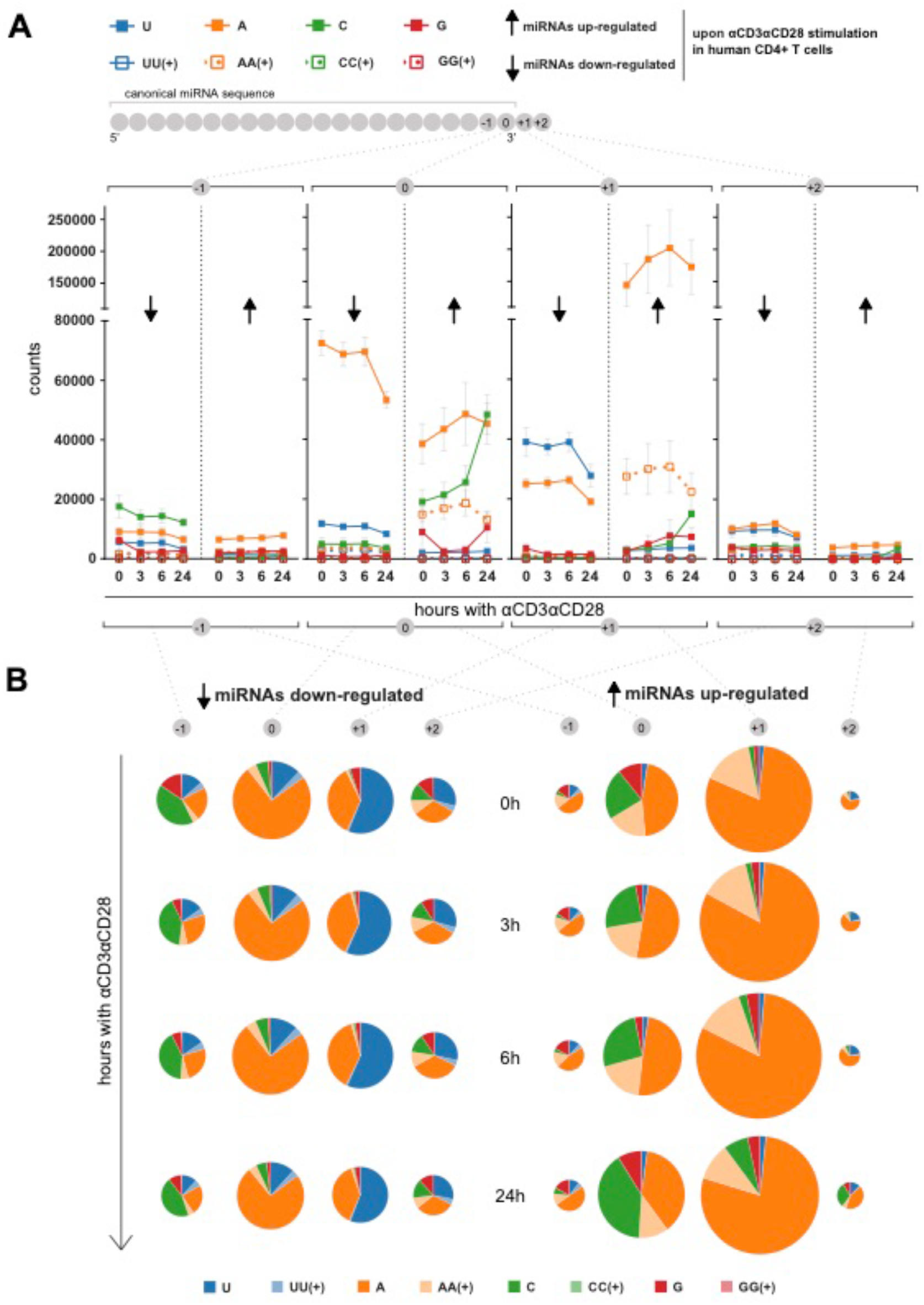
Kinetics of miRNA post-transcriptional modifications for αCD3αCD28 differentially expressed miRNAs. Kinetics of most abundant PtMs (mono-additions: U, A, C, G; and oligo-additions: ≥UU, ≥AA, ≥CC, ≥GG) at positions ‘3p-end −1’ (−1), ‘3p-end’ (0), ‘3p-end + 1’ (+1) and ‘3p-end + 2’ (+2); for up-regulated miRNA (left) and down-regulated miRNA (right) with an adjusted p-value <0.1. Mean and SEM (from three independent experiments) were plotted for each modification counts at specific positions across time points (A) and as pie charts whose area is proportional to the total number of counts for the specific position at indicated time points (B).

**Figure 5.**
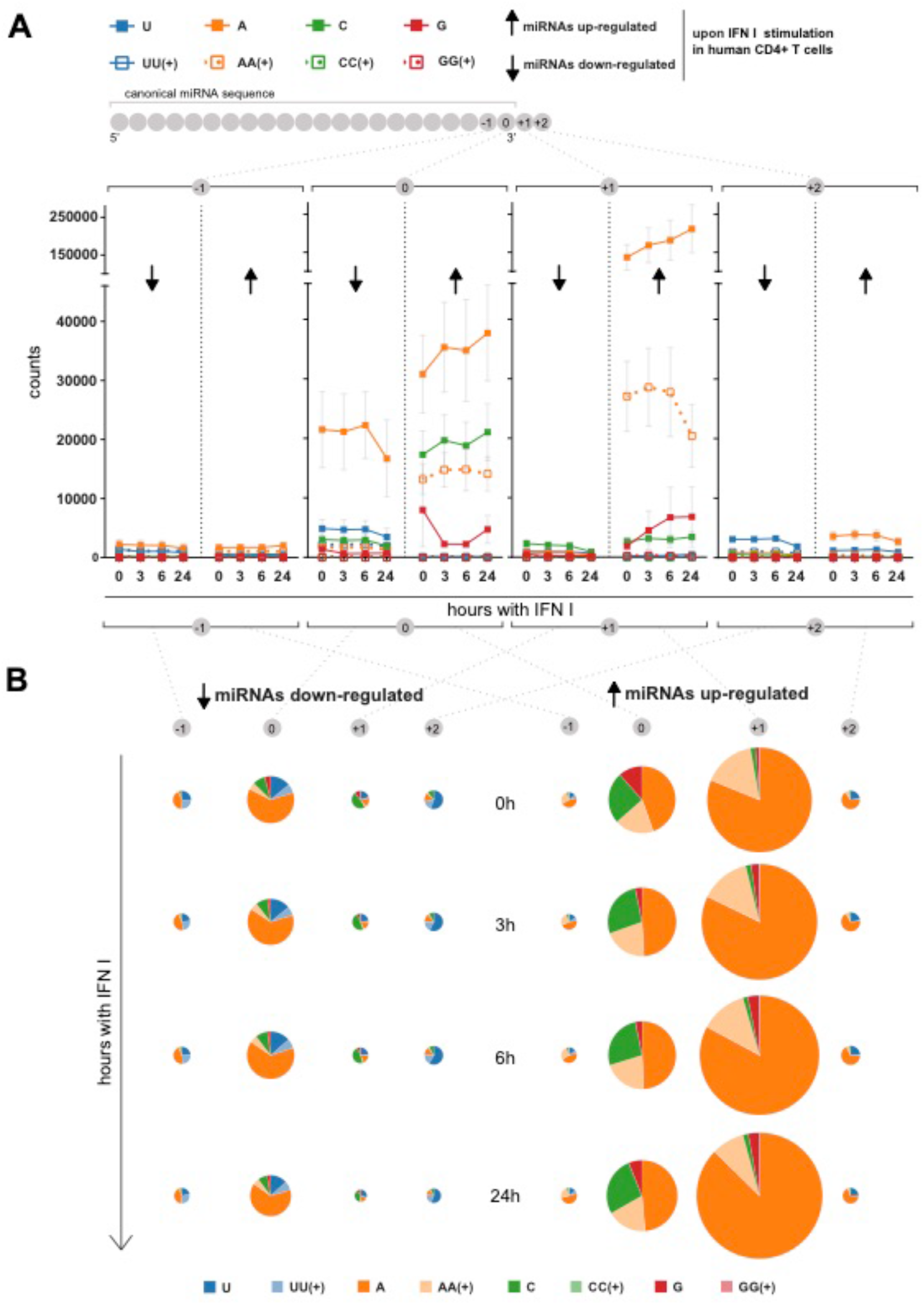
Kinetics of miRNA post-transcriptional modifications for IFN I differentially expressed miRNAs. Kinetics of most abundant PtMs (mono-additions: U, A, C, G; and oligo-additions: ≥UU, ≥AA, ≥CC, ≥GG) at positions ‘3p-end’ (0) and ‘3p-end + 1’ (1) and ‘3p-end + 2’ (2), for up-regulated miRNA (left) and down-regulated miRNA (right) with an adjusted p-value <0.1. Mean and SEM (from three independent experiments) were plotted for each modification counts at specific positions across time points (A) and as pie charts whose area is proportional to the total number of counts for the specific position at indicated time points (B).

### 4. Dis3l2, Eri1, TUT4 and TUT7 regulation upon T-cell activation

In addition, we evaluated the expression kinetics of four proteins related to RNA metabolism: TUT4 and TUT7 (terminal uridylyl transferases), and Dis3L2 and Eri1 (exonucleases that preferentially degrade uridylated RNA (Ustianenko et al. 2013; Chang et al. 2013; Hoefig et al. 2013)). For these experiments, we stimulated human CD4+ T cells from seven donors with αCD3αCD28 during various times up to 48 hours. A significant average upregulation was found after activation for all evaluated enzymes [Fig. 6A, 6B, 6C, 6D]. Early upregulation of Eri1 and Dis3L2 could be driving global miRNA downregulation upon T-cell activation. The overexpression of TUT4 and TUT7, could be indicating a higher uridylation activity, which could mark miRNA for degradation by Eri1 and Dis3l2.

**Figure 6.**
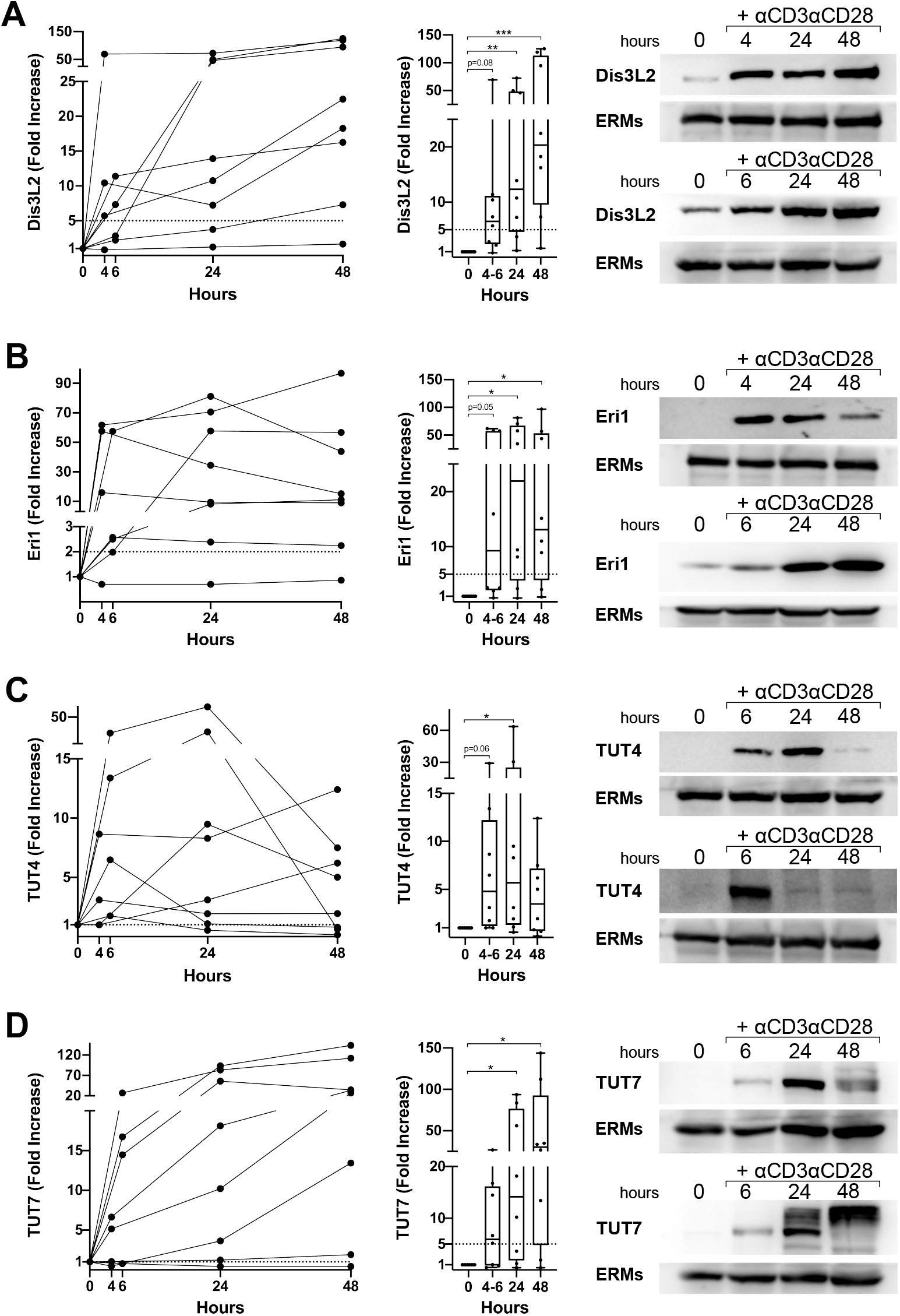
Expression of uridylated RNA degrading enzymes (Dis3L2 and Eri1) and terminal uridyl transferases (TUT4 y TUT7) upon CD4+ T cell activation. Western blot analysis of protein expression in human primary CD4+ T cell stimulated with αCD3αCD28, assessing Dis3L2 (A), Eri1 (B), TUT4 (C) and TUT7 (D). Fold increase compared to non-stimulation was represented for each donor to observe evolution upon activation (left panel) and using group median and interquartile range with whiskers ranging from minimum to maximum values (middle panel). Right panels include two examples for each protein to highlight inter-donor variability in upregulation kinetics. Band intensities were normalized to ERMs values and relativized to unstimulated conditions. Statistical analysis: Kruskal-Wallis test, Dunn’s multiple comparisons test [* p-value <0.05, ** p-value<0.01, *** p-value<0.001; 0.05<p-value<0.1: indicated with numbers]

## DISCUSSION

This study aimed to reveal the landscape of post-transcriptional modifications that could control miRNA levels during T cell activation with αCD3αCD28 and IFN I. We used NGS, since this technique allows the detection of isomiRs variants, offering also for the first time an unbiased exploration of miRNA differential expression in early time points of T cell activation.

Many miRNAs not previously linked with T cell stimulation were significantly down or up-regulated in response to αCD3αCD28. Two miRNAs could be highlighted for their differentiated behaviour: miRNA-1281 showed a high upregulation at early time points (3-6 h), and miRNA-4455 reached at 24 h a 122-fold increase, while no other upregulated miRNA surpassed 24-fold [Supplementary Fig. 2C]. MiRNA-1281 suppresses proliferation and migration in pulmonary artery smooth muscle cells, likely through targeting histone deacetylase 4 (HDAC4) (Li et al. 2018b). In osteosarcoma, miR-1281 (induced by the tumor suppressor p53) promoted apoptosis (Jiang et al. 2018). MiR-4455 also has a tumor suppressive effect, described in gastric cancer probably due to inhibition of ADGRD1 (Adhesion G protein-coupled receptor D1) (Zhou et al. 2019). ADGRD1 appears upregulated in a mouse model of gastrointestinal stromal tumor (Gromova et al. 2009) and inversely correlates with patient survival in acute myeloid leukemia (Yang et al. 2019) and glioblastoma (Bayin et al. 2016). It also plays an essential role in glioblastoma growth (Bayin et al. 2016; Frenster et al. 2017). Consistently, miR-4455 increased apoptosis and reduced proliferation, invasion and migration in a gastric cancer cell line (MGC-803) (Chen et al. 2018). MiR-4455 targets VASP 3’-UTR and a decrease in VASP levels results in PI3K/AKT inhibition (Chen et al. 2018). VASP is a key mediator of actin cytoskeleton remodeling, polarization and cell migration (Krause et al. 2003; Bailly 2004). Ena/VASP proteins are part of a complex that connects signaling from TCR receptor to actin remodeling (Krause et al. 2000). A previous study used a double knock out mouse to demonstrate that EVL (Ena-VASP-like) and VASP are involved in activated T-cell trafficking into inflammation sites and secondary lymphoid tissues, particularly promoting diapedesis (Estin et al. 2017). In light of these effects, the strong effort of T cells for up-regulating miR-1281 and miR-4455, respectively, earlier and higher than any other miRNA, might be related with generating a negative feedback for the control of T cell activation.

To our knowledge, no other study has evaluated miRNA changes in human primary T cells stimulated with type I IFN. A 2015 review gathered data available on IFN I regulated microRNAs, mainly in the liver cell line Huh7 and human glioma (Forster et al. 2015). Out of 36 miRNAs identified, 7 were described in two or more studies, indicating certain overlapping but also a great diversity across cell types regarding their response to IFN I (Forster et al. 2015). In fact, only one of these 36 miRNAs (miR-212) has been found as differentially expressed in our samples. Two prior studies had used immune cells: PBMCs and NK cells. MiRNAs involved in the anti-viral response against Hepatitis C virus (miR-1, miR-30, miR-128, miR-196 and miR-296) were induced in peripheral blood mononuclear cells (PBMCs) upon IFN-α treatment (Scagnolari et al. 2010). In human NK Cells, miRNA-30e and miRNA-378 were downregulated by IFN I (Wang et al. 2012). Our study provides a novel dataset of IFN I regulated miRNAs in human primary CD4 T cells, which comprises 24 miRNAs that are upregulated and 33 that are downregulated. Of the 57 genes modulated by IFN-I, 37 are also regulated by αCD3αCD28 stimulation. Interestingly, many miRNAs significantly upregulated in our samples, such as miR1246(Qi et al. 2013; Li et al. 2017), miR-1261 (Martins et al. 2018), miR-1290 (Qi et al. 2013), miR-3196(Herkt et al. 2020), miR-3614-5p(Diosa-Toro et al. 2017), miR-4301(Diosa-Toro et al. 2017), miR-4448 (Baños-Lara et al. 2018) or miR-4488 (Alipoor et al. 2017), also appear increased in cells infected with Dengue, RSV (Respiratory Syncytial Virus), Hepatitis C virus (HCV), Legionella pneumophila or BCG (Mycobacterium bovis Bacillus Calmette–Guerin).

IPA assessment of IFN-I and αCD3αCD28 regulated miRNAs indicates their involvement in cellular development, growth, proliferation and movement. These processes are indeed essential for activated T cells to perform their function, which includes their differentiation to effector and/or memory phenotypes to combat infection short- and long-term, respectively. In this regard, activated T cells undergo intense cellular reprogramming with an increase in mRNA and protein expression. Accordingly, activated T cells would need to control inhibitory safeguards that prevent abnormal activation that could produce autoimmunity. MiRNA regulation may act as a negative regulator of gene expression, which would need to be withdrawn, at least partially. Several studies support this hypothesis. For instance, mRNAs undergo 3’ UTR shortening upon T lymphocyte activation, thereby reducing the pool of potential target sites for miRNA binding (Sandberg et al. 2008). Moreover, T cell activation promotes a rapid global miRNA downregulation and degradation of Argonaute proteins, which are key effectors of the RISC complex (Bronevetsky et al. 2013). We hypothesize that an active mechanism of miRNA degradation underlies the intense miRNA downregulation observed only a few hours after T cell activation. For this reason, we evaluated the expression of Eri1 and Dis3L2. Both exoribonucleases display a clear preference for uridylated RNA substrates (Ustianenko et al. 2013; Chang et al. 2013; Hoefig et al. 2013); also, Eri1-deficient NK cells and T cells showed increased overall miRNAs levels (Thomas et al. 2012). Here we detected a marked upregulation of both enzymes following T cell activation. Consistently, TUT4 and TUT7, which are specifically regulated by αCD3αCD28 stimulation, could be uridylating substrates and labelling them for subsequent degradation by Eri1 or Dis3l2.

Nevertheless, T cells still require certain miRNAs to remain stable even in generally degradative conditions. PtMs could control miRNA stability, promoting degradation but also protecting specific miRNAs. The global PtMs profile of our samples reveals that modification processes were focused on the 3’ end of most miRNAs. While uridylation and adenylation have been the best characterized 3’ end modifications described across animal miRNAs (Landgraf et al. 2007; Chiang et al. 2010; Burroughs et al. 2010; Wyman et al. 2011; Muller et al. 2014), cytosylation was highly represented in our samples. Cytosine was specifically found at position 0 (3’ end nucleotide). At this position, A is equally frequent and much more represented than U and G. Unexpectedly, upregulated miRNAs displayed increased levels of C addition compared to those downregulated under the same conditions. Upregulated miRNAs were also characterized by a strong presence of mono-A at 1a and oligo-A at 0 and 1. Consistent with previous studies from our laboratory, which had indicated that urydilation is a miRNA degradation signal in T cells (Gutiérrez-Vázquez et al. 2017), higher levels of U additions were found in αCD3αCD28 down regulated miRNAs. A similar pattern can be observed in IFN I stimulation, although differences are milder, which may be due to a less dynamic miRNA environment; the number of differentially expressed miRNAs was roughly half of those quantified in cells treated with αCD3αCD28.

Although most studies evaluating PtMs have found guanosine and cytosine additions to be barely represented, mono-addition of cytosine was the second most abundant 3’ modification after mono-uridylation, in mouse primordial germ cells and gonadal somatic cells at various embryonic stages (Darnell et al. 2018). The presence of ‘non-templated cytosylation’ has been described in Arabidopsis, which prompted the hypothesis of the existence of a nucleotidyl transferase with a preference for cytosine as substrate (Chou et al. 2015). Cytosine additions could be relevant for miRNA in very specific developmental or differentiation stages.

In summary, this study offers a novel dataset of differentially regulated miRNAs in early time points of human primary CD4+ T cell activation. Most importantly, we also describe the kinetics of post-transcriptional modifications and the potential effect of these modifications in miRNA stability in the context of T cell activation. Our data also indicate that RNA degrading enzymes Eri1 and Dis3L2 are upregulated upon activation, which could be part of an active mechanism of miRNA degradation guided by uridylation. Indeed, higher uridylation was found in downregulated miRNAs. Unexpectedly, upregulated miRNAs, which manage to multiply their levels in this adverse environment, point towards 3’ cytosine and adenine additions as a protective signal.

## METHODS

### 1. Human primary CD4 T cell culture

Human peripheral blood mononuclear cells (PBMCs) were isolated from buffy coats, obtained from healthy donors, by separation on Biocoll Separating Solution (Biochrom, L6115) according to standard procedures. Non-adherent cells were separated from PBMCs after a 30 min adherence step at 37°C. CD4^+^ T cells were purified from non-adherent cells using Human Resting CD4+ T cell Isolation Kit (STEMCELL Technologies, 17962). A specific reagent to isolate resting T cells was selected to avoid the presence of pre-activated CD4+ T cells in sequencing samples. In experiments performed to evaluate protein expression, CD4+ T cells were isolated with EasySep Human CD4 T Cell Isolation Kit (STEMCELL Technologies, 17952).

For T cell stimulation, we treated CD4+ T cells with either αCD3αCD28 (ImmunoCult™ Human CD3/CD28 T Cell Activator; STEMCELL Technologies, 10971) or IFN I (1:1000, Human IFN Alpha Hybrid (Universal Type I IFN); PBL ASSAY SCIENCE, 11200-1). Cells were cultured in RPMI 1640 (Gibco), supplemented with 10% fetal bovine serum (Sigma), 20mM Hepes (Hyclone), 0.3mg/mL L-glutamine (Hyclone), 100 U/mL penicillin (Gibco) and 100 μg/mL streptomycin (Gibco).

These studies were performed according to the principles of the Declaration of Helsinki and approved by the local Ethics Committee for Basic Research at the Hospital La Princesa (Madrid); informed consent was obtained from all human volunteers.

### 2. RNA isolation, library preparation and NGS

Three independent experiments, with resting CD4+ T cells isolated from different healthy donors were performed. Samples were collected at 0h and after αCD3αCD28 or IFN I stimulation during 3h, 6h and 24h. The 21 samples were lysed in QIAzol Lysis Reagent (Qiagen, 79306) and RNA was extracted using the miRNeasy Mini Kit (Qiagen, 217004). In order to reduce phenol-based reagent contaminations, purified RNA samples were precipitated using sodium acetate (3M, 0.1x sample volume) and ethanol (100%,3x sample volume). RNA integrity was evaluated using an Agilent 2100 Bioanalyzer (Eukaryote Total RNA Nano assay).

A total of 200 ng of total RNA were used to generate barcoded miRNA-seq libraries using the NEBNext Multiplex SmallRNA Library Prep Set for Illumina (New England Biolabs). Briefly, 3’ and 5’ SR adapters were first ligated to the RNA sample. Next, reverse transcription followed by PCR amplification was used to enrich cDNA fragments with adapters at both ends. The quantity and quality of the miRNA libraries were determined using the Agilent 2100 Bioanalyzer High Sensitivity DNA chip.

Libraries were sequenced on a HiSeq2500 (Illumina) to generate 60 bases single reads. FastQ files for each sample were obtained using bcltofastQ 2.20 Software software (Illumina). NGS experiments were performed in the Genomics Unit of the CNIC.

### 3. miRNA-Seq data analysis

Sequencing reads were pre-processed by means of a pipeline that used FastQC (http://www.bioinformatics.babraham.ac.uk/projects/fastqc/), to asses read quality, and Cutadapt (Martin 2011) to trim sequencing reads, eliminating Illumina adaptor remains, and to discard those that were shorter than 15 nt or longer than 35 nt after trimming. Around 80% of the reads from any of the samples were retained. Resulting reads were aligned against a collection of 2657 human, mature miRNA sequences extracted from miRBase (release 22), to obtain expression estimates with RSEM (Li and Dewey 2011). Percentages of reads participating in at least one reported alignment were around 40%. Expected expression counts were then processed with an analysis pipeline that used Bioconductor package Limma (Ritchie et al. 2015) for normalization (using TMM method) and differential expression testing, taking into acount that samples had been obtained in three batches, and considering only 626 miRNA species for which expression was at least 1 count per million (CPM) in 3 samples. Changes in gene expression were considered significant if associated to Benjamini-Hochberg adjusted p-value < 0.1. Clustering of expression profiles and production of heatmaps were performed with Genesis (Sturn et al. 2002). Epi-transcriptomic modifications were detected with Chimira (Vitsios and Enright 2015), an online tool that, after alignment of miRNA-Seq reads against miRBAse records, identifies mismatched positions to classify them and to quantify multiple types of 3'-modifications (uridylation, for example), as well as 5'-modifications and internal modifications or variations. Count tables produced by Chimira were further processed with ad-hoc produced R-scripts to normalize modification counts by library size and to calculate summary statistics across groups of replicate samples. Analyses were restricted to the collection of 626 miRNA species with detectable expression.

Two core analysis were performed by Ingenuity Pathway Analysis (Content version: 49932394 (Release Date: 2019-11-14), one with all miRNAs differentially expressed at least one time point of stimulation with αCD3αCD28 and a second one with the corresponding IFN-I regulated miRNAs. MiRNA-target networks were buildt with miRNet loading the highest upregulated and downregulated miRNAs for each treatment (Fan and Xia 2018). Ven diagrams were elaborated with Venny (https://bioinfogp.cnb.csic.es/tools/venny/index.html).

### 4. Immunoblotting

Cells extracts were prepared in lysis buffer (50 mM Tris pH 7.5, 150 mM NaCl, 1%NP-40, 5 mM EDTA, 50mM NaF, 5mM DTT) supplemented with a protease inhibitor cocktail (Complete, Roche). Cell lysates were cleared of nuclei by centrifugation (15000 g, 15 min). Proteins were separated on 8-10% SDS-PAGE gels and transferred to a nitrocellulose membrane. Membranes were incubated with primary specific antibodies: anti-Dis3L2 (Nobus biologicals, NBP1-84740), anti-Eri1/THEX1 (Cell Signaling, #4049), anti-TUT4/ZCCHC11 (Pro Sci incorporated, 46-610), anti-TUT7/ZCCHC6 (Proteintech, 25196-1-AP), and anti ezrin/moesin (ERMs) (90/3) (provided by Heinz Furthmayr, Stanford University, CA). Primary antibodies were used at 1:2000 dilution and peroxidase-conjugated secondary antibodies (goat anti-rabbit, ThermoFisher Scientific #31460; rabbit anti-goat, Thermo Scientific #31402) at 1:5000. Chemoluminescence was measured with LAS-3000 (Fujifilm). Band intensities were quantified with Image Studio Lite (LI-COR Biosciences), normalized to ERMs values and relativized to unstimulated conditions (when no band was detected at 0 h, background was taken as reference signal).

## DATA ACCESS

### GEO SUBMISSION

All raw and processed sequencing data generated in this study have been submitted to the NCBI Gene Expression Omnibus (GEO; https://www.ncbi.nlm.nih.gov/geo/) under accession number GSE156287. Data are currently in private status but can be reviewed by entering token utahemcclrkxzkt into the corresponding box.

### UCSC GENOME BROWSER SESSIONS

Alignments are accessible at the following UCSC Genome Browser session:

* https://genome.ucsc.edu/s/mjgommo/ARG_CD4T_IFN_I
* https://genome.ucsc.edu/s/mjgommo/ARG_CD4T_aCD3aCD28

Each session has been configured to allow the visualization of 13 custom tracks, which consist in:

- miRBase_mature track: representing the coordinates of all mature miRNAs described in miRBase, release 22.
- 12 BAM alignment tracks, corresponding to three replicate samples for the control condition (0h) and each of the time points (3h, 6h, 24h) for IFN I or αCD3αCD28 treatment.

As previously described, miRNA detection and quantification has been performed in this study by aligning NGS processed reads against a transcriptomic reference consisting in all mature miRNA sequences described in miRBAse, release 22, for Homo sapiens. To produce genomic alignments that were fully congruent with those generated for quantification, BAM alignments displayed in the UCSC tracks have been generated with RSEM using a genomic reference constructed with the mature miRNA coordinates described in miRBase, release 22, exclusively. For this reason, coverage is expected only on the intervals corresponding to regions that code for mature miRNAs. MIMAT IDs are used to identify such intervals because they are guaranteed to be unique (locus specific). Visualization of the tracks may require reloading the page, because of timeout issues.

### IN-HOUSE SCRIPTS

Chimira results describing miRNA modifications consist in a separated table for each sample. A specialized, in-house R script (CHIMProcessor.R) was developed to process the collection of output files, as well as several other auxiliary files, in order to normalize modification frequencies by library size, merge frequency information from replicate samples, filter data using various parameters and generate combined tables and preliminary plots. The script is available from GitHub, at:

* https://github.com/mjgommo/CHIMProcessor

together with Chimira modification output files produced in the context of the current study, as well as auxiliary files required to reproduce the presented results.

## ACKNOWLEDGMENTS

We thank Miguel Vicente-Manzanares for help with English. We also thank the CNIC Genomics and Bioinformatics Units for technical support.

This work was supported by grants SAF2017-82886-R from the Spanish Ministry of Economy and Competitiveness (MINECO), grant S2017/BMD-3671-INFLAMUNE-CM from the Comunidad de Madrid, a grant from the Ramón Areces Foundation “Ciencias de la Vida y la Salud” (XIX Concurso-2018) and a grant from Ayudas Fundación BBVA a Equipos de Investigación Científica (BIOMEDICINA-2018), the Fundació Marató TV3 (grant 122/C/2015) and “La Caixa” Banking Foundation (HR17-00016). BIOIMID (PIE13/041) from Instituto de Salud Carlos III, CIBER Cardiovascular (CB16/11/00272, Fondo de Investigación Sanitaria del Instituto de Salud Carlos III and co-funding by Fondo Europeo de Desarrollo Regional FEDER). The Centro Nacional de Investigaciones Cardiovasculares (CNIC, Spain) is supported by the Ministerio de Economía y Competitividad-Spain and the Pro-CNIC Foundation.

ARG, SGD and IFD were supported by the FPU program (Spanish Ministry of Education).

## AUTHOR CONTRIBUTIONS

ARG and FSM conceived the study. Experiments were performed by ARG, SGD and IFD. MJG and FSC performed the differential expression and post-trancriptional modifications analysis of the small RNAseq data. ARG analysed results, made the figures and wrote the manuscript, with input from the rest of authors. FSM supervised and revised all the work.

